# A Robust Expression and Purification Protocol for the Production of the La Domain of Human LARP6

**DOI:** 10.1101/2024.06.11.598414

**Authors:** Blaine Gordon, Nolan Blackford, Robert Silvers

## Abstract

Human La-related protein 6 (HsLARP6) regulates the highly organized biosynthesis of type I procollagen polypeptides and affects proper assembly of procollagen peptides into heterotrimers of type I procollagen. HsLARP6-mediated regulation of collagen biosynthesis is mediated through interaction with the 5’ stem loop (5’SL) motif found in type I and III collagen messenger RNA. Recent studies highlight the involvement of HsLARP6 in fibroproliferative diseases and its potential as a target for therapeutic intervention. The intrinsic propensity of the La domain of HsLARP6 to aggregate hampers studies probing the molecular basis of biologically- and disease-relevant structure-function relationship, particularly when high concentrations are required. This work provides detailed procedures to produce milligram amounts of RNase-free and functional La domain of HsLARP6. Furthermore, we investigated the effect of the protein construct length and RNA binding on protein stability. C-terminal truncations greatly impact protein stability, while N-terminal truncations have little to none effect on protein aggregation and RNA binding. When in complex with its cognate 5’SL RNA, the La domain shows unprecedented stability compared to the aggregation-prone unbound state. The protein-RNA complex remains stable for at least 50x longer than the unbound state, under identical conditions. These results provide a significant platform for further studies of the molecular recognition of 5’SL by HsLARP6.

## Introduction

Human La-related protein 6 (HsLARP6) belongs to a diverse, but highly conserved superfamily of RNA-binding proteins (RBPs) that is involved in gene regulation at the post-transcriptional level.^1,2^ Due to their pivotal role in the regulation of the intricate balance of translation, breakdown, and storage of messenger RNAs (mRNAs), LARPs are closely linked to the onset and progression of various cancers, neurodegenerative disorders, and fibroproliferative diseases.^3–8^

The LARP6-mediated regulation of type I and III collagen biosynthesis plays a pivotal role in the onset and progression of fibroproliferative diseases.^7^ Type I collagen is predominantly produced by fibroblasts and myofibroblasts and is one of the major constituents of the extracellular matrix of tendons, skin and bones, while it is only present in small amounts in healthy tissues and organs such as liver, heart, kidney, and lungs. In fibroblasts, the type I collagen polypeptides α1(I) and α2(I) are translated in a highly regulated and organized process and the folding of type I procollagen heterotrimers within the endoplasmic reticulum occurs in organized structures termed collagenosomes.^8^ In fibroproliferative disease, the biosynthesis of type I procollagen peptides is significantly elevated in affected tissues, leading to scarring and impairment of tissue function.^9,10^ It was shown that LARP6 facilitates biosynthesis by recruiting accessory translational factors, which enhance the translational efficiency.^11–16^ The interaction between LARP6 and the collagen mRNA is directly linked to this increase in collagen peptide biosynthesis.^17^ Furthermore, it was recently shown that the interaction between LARP6 and collagen mRNAs is crucial for the proper formation of collagenosomes.^8,16,18,19^ Furthermore, in an animal model where the LARP6-dependent pathway of type I collagen biosynthesis is interrupted, it was shown that resulting animals are resistant to hepatic fibrosis, underlining the importance of LARP6’s interaction with its cognate RNA in fibrotic disease.^7^ These findings highlight the critical nature of the LARP6 interaction with collagen mRNA, that contributes to disease progression, providing potential targets for therapeutic intervention in fibrosis and other related conditions.

The defining structural feature of all members of the LARP superfamily is the presence of an RNA-binding domain (RBD), known as the La domain (Figure 1A). In the majority of LARPs, the La domain (LA) is paired with a downstream RNA recognition motif (RRM) connected by a short linker. Both domains form a functional unit, termed the La module (LAM), that is essential for mediating the interaction between the LARP and their target mRNAs.

**Figure 1:**
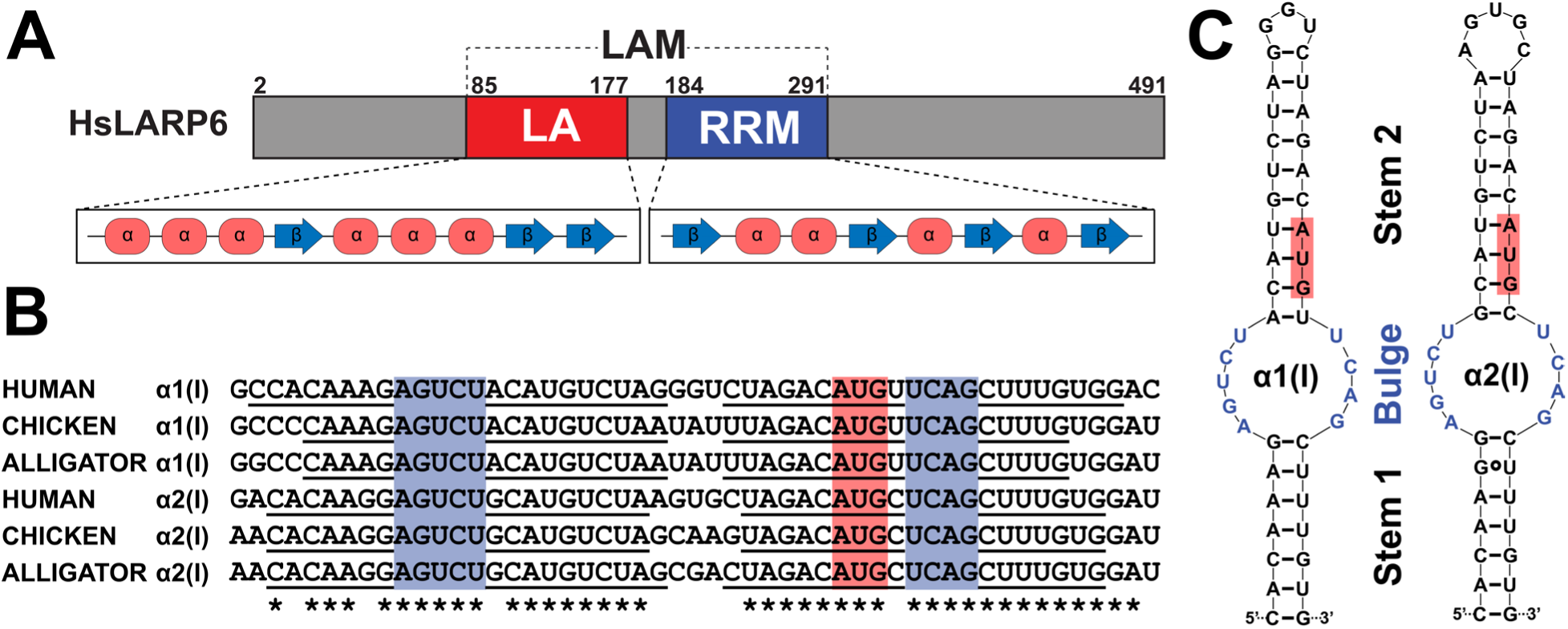
Architecture of HsLARP6 and 5’SL RNA. (A) Domain organization of HsLARP6 highlighting the La domain (LA) and RNA recognition motif (RRM) which compose the La module (LAM). Secondary structure of LA and RRM domains are as indicated. (B) Sequence alignment of mRNAs encoding the α1(I) and α2(I) strands of type I collagen from *H. sapiens*, *G. gallus*, and *A. mississippiensis*. Highly conserved stems (underlined) surround entirely conserved bulge regions (blue). Conserved bases are indicated by an asterisk (*). The location of the start codon is shown in red. (C) Secondary structure prediction for human α1(I) and α2(I) 5’SL. Bases belonging to the bulge and start codon are shown in blue and red, respectively.

Commonly, LARPs recognize single-stranded RNA motifs. LARP1 binds the poly(A) tail in mRNA as well as the 5’ terminal oligopyrimidine (5’TOP) motif in TOP mRNAs,^20–22^ LARPs 3 and 7 bind the UUU-3’OH motif of RNA polymerase III transcripts and 7SK snRNA, respectively,^23–25^ while LARP4 interacts with AU-rich sequences.^26^ LARP6 however recognizes a double-stranded RNA motif known as the 5’ stem-loop (5’SL, Figure 1B) that is highly conserved across all vertebrates and plays a critical role in regulating the translation of collagen mRNAs.^16,27–29^ The 46-48 nucleotides long 5’SL RNA is located at the junction between the 5’ untranslated region (5’UTR) and coding region of mRNAs coding for the type I collagen peptides α1(I) and α2(I) as well as type III collagen peptide α1(III). Secondary structure prediction (Figure 1C) of this region indicates that the 5’SL forms a stem loop structure containing an internal loop (bulge) harboring nine highly conserved nucleotides flanked by two stems. The AUG start codon is located within stem 2 of the 5’SL motif.

The structure of a complex between HsLARP6 and 5’SL is currently unknown; hence the molecular basis of the interaction remains poorly understood. Progress is further hampered by conflicting studies regarding the individual roles of the La domain and RRM in 5’SL binding (*vide infra*). Here, we report an improved strategy for recombinant production of milligram quantities of high-purity La domain that is RNase-free and binding-competent, a prerequisite for biochemical and biophysical studies of 5’SL binding. We show that the choice of protein construct modulates the stability and binding competency of the La domain by probing N- and C-terminal extensions of different lengths. Furthermore, we show that the inherent instability of the La domain is suppressed entirely by the binding of its cognate 5’SL RNA. These results provide insights into the binding of La domain to the 5’SL while also providing a significant platform for further studies into the molecular basis of binding.

## Results and Discussion

The post-translational regulation of collagen biosynthesis mediated by HsLARP6 and its role in the onset and progression of fibroproliferative diseases is well documented. The interaction of the La module and 5’SL is directly involved in this process,^16,18,19^ hence several previous studies have focused on studying the interaction employing biochemical and biophysical methods.

Access to a detailed molecular understanding of 5’SL RNA binding to the La module of HsLARP6 has been hampered by various factors. To begin with, the La domain and La module of HsLARP6 are intrinsically unstable and aggregation-prone,^16,27^ which has also been observed for LARPs from other species including XmLARP6 and DrLARP6a,^30^ complicating production, handling, and storage of protein preparations. Secondly, the La domain of HsLARP6 has a basic surface unique among human LARPs, making it prone to non-specific interactions with lingering nucleic acids during purification that interfere with downstream binding assays. Finally, we show that the La domain of HsLARP6 is prone to co-purification with host RNases.

Below, we describe a detailed roadmap to obtaining milligram quantities of RNase-free, high-purity HsLARP6 La domain, a prerequisite for further studies into the binding of the La domain to its cognate 5’SL RNA. Additionally, we describe the dependency of La domain stability and binding competency on the length of N- and C-terminal extension. Finally, we show that the formation of the protein-RNA-complex suppresses the instability of the La domain entirely.

### Preparation of RNase-free, high-purity La domain

The preparation of high-purity and RNase-free La domain of HsLARP6 is a prerequisite for any investigations into the binding of 5’SL to the La domain and La module of HsLARP6, in particular where high working concentrations are required. Previous *in vitro* purifications of the HsLARP6 La domain have either been single step immobilized metal affinity chromatography (IMAC) purifications^16^ or resulted in binding measurements inconsistent with findings from two separate sources.^27,31^ Furthermore, the previously published protocols for protein expression and purification resulted in inconsistent results in our hands with protein preparations still containing host RNases and nucleic acid impurities that impeded biochemical and biophysical studies. Our studies also showed that host ribonucleases (RNases) tend to co-purify with the La domain of HsLARP6.

We recombinantly expressed various La domain constructs in *E. coli* and purified them using multiple steps of affinity and size exclusion chromatography steps. The constructs used here vary in length of their N- and C-terminal extension and are named HsLARP6(73-183), HsLARP6(79-183), HsLARP6(84-183), HsLARP6(73-176), and HsLARP6(73-171). The same expression and purification protocol was followed for all the aforementioned constructs. In the following, we focus on the detailed description of the RNase-free production of homogenous, monodisperse HsLARP6(79-183).

Using sodium-dodecyl-sulfate polyacrylamide electrophoresis (SDS-PAGE), the production and purity of HsLARP6(79-183) at various purification steps was analyzed (Figure 2A). Simultaneously, the presence of lingering host RNases during purification was determined by RNase activity assay (Figure 2B). After the cell lysate was cleared from insoluble debris, a strong overexpression band of His_6_-TEV-HsLARP6(79-183) can be seen (Figure 2A, lane 1) and the cell lysate expectedly shows RNase activity (Figure 2B, lane 1). During the IMAC purification, two separate wash steps are involved. The first wash is done at low salt concentration, the second wash is performed using a high-salt buffer. RNase activity in the high-salt wash indicates that lingering host RNases cannot be removed with the low-salt wash alone (Figure 2B, lane 2). After elution of His_6_-TEV-HsLARP6(79-183) from the IMAC column, the protein is already quite pure (Figure 2A, lane 3), and more importantly, the elution does not contain any appreciable amounts of RNase (Figure 2B, lane 3). The high-salt wash is mandatory to remove any lingering host RNases. Interestingly, the intact mock RNA does not penetrate the gel when mixed with the fraction collected after elution from the IMAC (Figure 2B, lane 3), likely due to large complex formation with eluate components, which does not affect our conclusion that the elution fraction after IMAC is RNase-free. Complex formation of the mock RNA is not observed after His_6_-tag removal by TEV digestion or beyond. High purity and RNase-free HsLARP6(79-183) is obtained after reverse IMAC to remove His_6_-tagged TEV protease (Figures 2A & B, lane 4), as well as subsequent ion exchange chromatography (Figures 2A & B, lane 5) and size exclusion chromatography (Figures 2A & B, lane 6).

**Figure 2:**
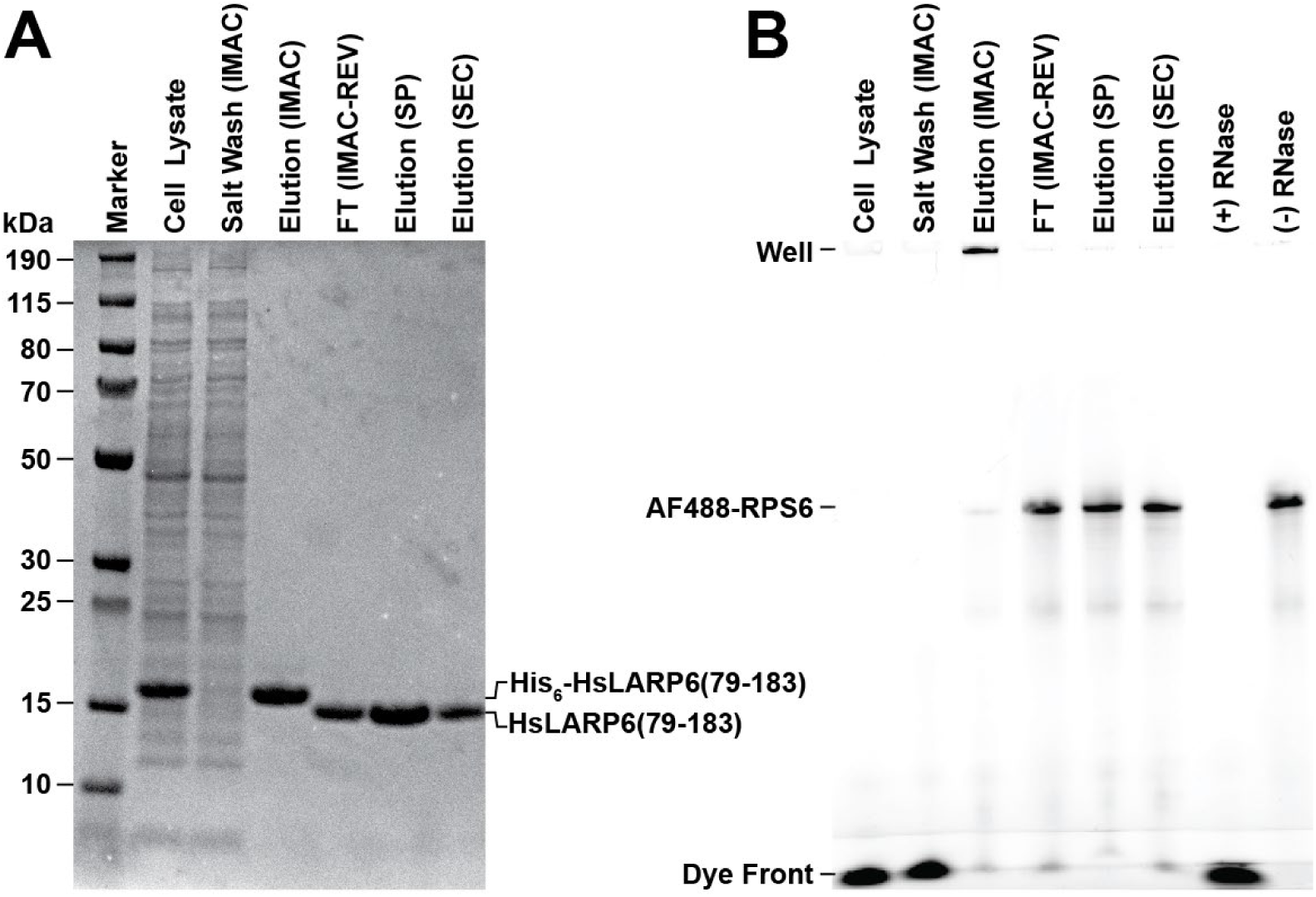
Expression and RNase-free purification of HsLARP6(79-183). (A) SDS-PAGE of several purification steps. Left to right: PageRuler™ Plus prestained protein ladder (marker), cell culture lysate supernatant, pooled fractions from the high salt (1M NaCl) wash performed in the IMAC purification step, elution from the first IMAC, flowthrough (FT) of the reverse IMAC after TEV digestion, elution from the SP cation exchange step, and elution from SEC. The positions of each protein species before and after TEV digestion are labeled. (B) RNase activity assay. Fluorescence image of 8M urea denaturing PAGE of the same samples from each purification step shown in A incubated at 37°C in the presence of a fluorescent RNA (AF488-RPS6 41mer) followed by a positive control containing 0.1 mg/mL RNase A and a negative control. The position of the well, AF488-RSP6 41mer, and dye front are labeled.

From the expression in 1 L LB medium, a 4.5 g wet cell pellet yielded a total of ~25 mg of high-purity, RNase-free HsLARP6(79-183). After size exclusion chromatography, an A_260_/A_280_ ratio of 0.54 was obtained, indicating a complete to near-complete removal of nucleic acids contaminants.^32^

### The La domain exhibits high-affinity binding to 5’SL RNA

It was previously suggested that the La domain of HsLARP6 is unable to bind the 5’SL RNA and that the full-length La module including the downstream RRM domain is required for binding.^16,27^ Isothermal Titration Calorimetry (ITC) data of HsLARP6(70-183) presented alongside its solution NMR structure by the Conte laboratory indicated no binding between the La domain and 5’SL.^27^ In contrast, we recently showed in collaboration with the Stefanovic laboratory that the La domain itself is sufficient for binding 5’SL RNA.^31^ Indeed, binding assays performed on freshly prepared La domain and 5’SL using microscale thermophoresis (MST) and NMR spectroscopy showed high-affinity binding to 5’SL in the absence of the downstream RRM domain with great reproducibility (Figure 3).

**Figure 3:**
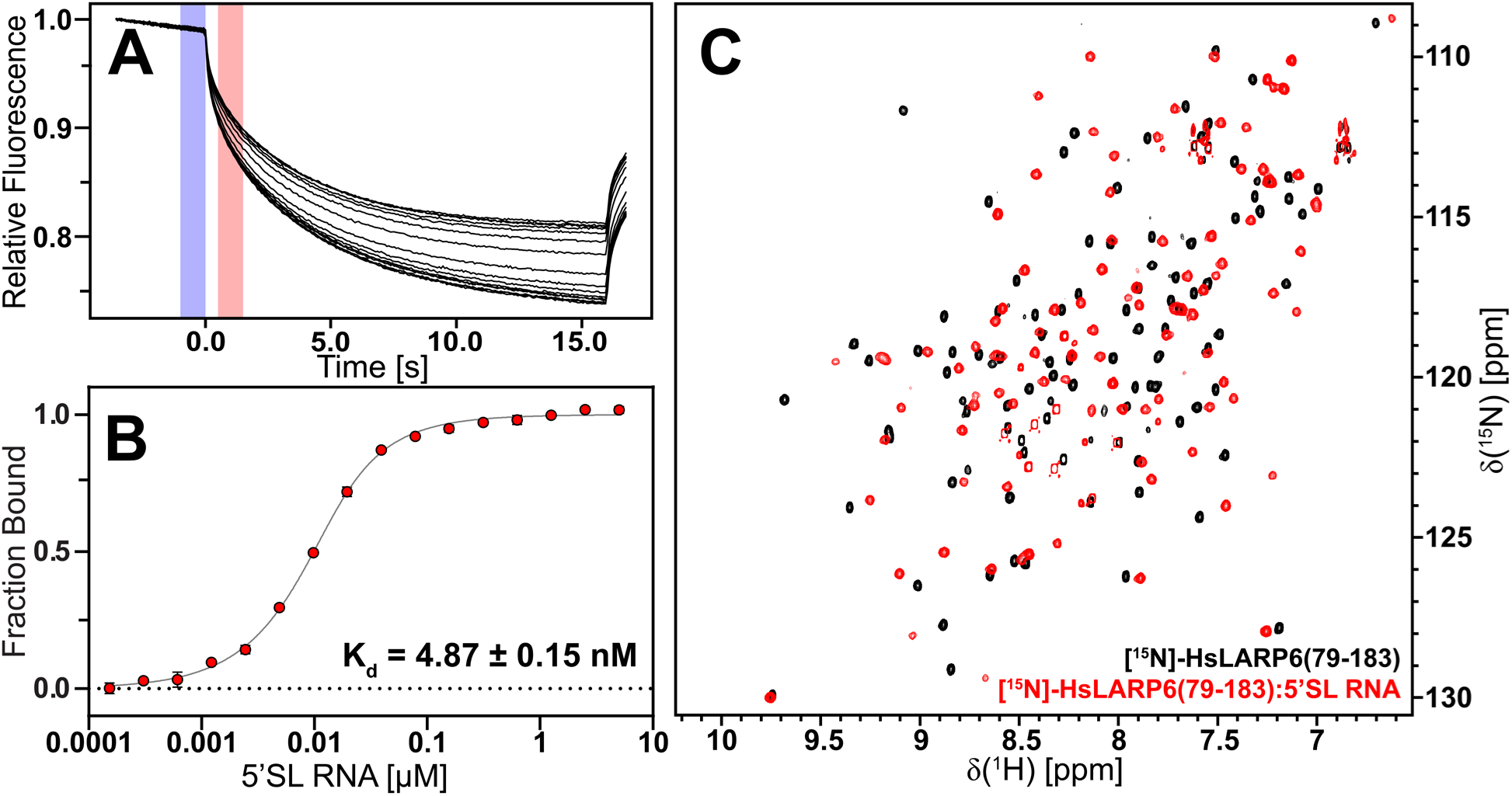
The La domain of HsLARP6 binds to 5’SL RNA with high affinity. (A) Microscale thermophoresis traces (black) from each of the 16 titration points used to determine the equilibrium dissociation constant between Cy5-LARP6(73-183)G74C and 5’SL RNA at 25 °C. “Cold” and “Hot” regions are shown in blue and red, respectively. (B) Fitted binding curve from microscale thermophoresis of Cy5-HsLARP6(73-183)G74C against 5’SL RNA at 25 °C. The dissociation constant K_D_ is shown. (C) [^1^H,^15^N]-HSQC spectra of [^15^N]-HsLARP6(79-183) unbound (black) and bound (red) to 5’SL RNA. Spectra were collected at 298 K and 16.4 T.

Microscale thermophoresis performed on Cy5-labeled La domain demonstrated the ability of the La domain to bind 5’SL with a K_D_ of 4.87 ± 0.15 nM (Figure 3A&B). This is in agreement with previously reported K_D_ values ranging from 0.5 to 40 nM for the binding between various HsLARP6 species and 5’SL RNA.^19,27,31^ Similarly, [^1^H, ^15^N]-HSQC spectra of isotopically-labelled unbound and 5’SL-bound La domain samples showed clear chemical shift perturbations (CSPs) which evidence the formation of the 1:1 complex (Figure 3C). The absence of any resonances stemming from the unbound state in the spectrum of the bound sample indicated the complete conversion of the unbound state into the bound state and is indicative of high-affinity binding. Furthermore, analytical size exclusion chromatography (SEC) performed on La domain constructs of varying lengths showed a full conversion of the unbound state to the bound state (*vide infra*, Figure 6).

### N- and C-terminal extensions modulate stability and 5’SL binding

The previously published NMR structure of the unbound La domain of HsLARP6 suggested that the folded core domain spans residues 85 to 171 (Figure 4A).^27^ Upstream of residue 85 and downstream of residue 171, the average pairwise RMSD of C^α^ atoms rises sharply (Figure 4B), indicating the N-terminal and C-terminal extensions are flexible. To examine how the presence of these extensions affects protein stability and binding competency, we investigated a total of five constructs, three probing the N-terminal extension and two probing the C-terminal extension beyond the folded protein core structure (Figure 4C).

**Figure 4:**
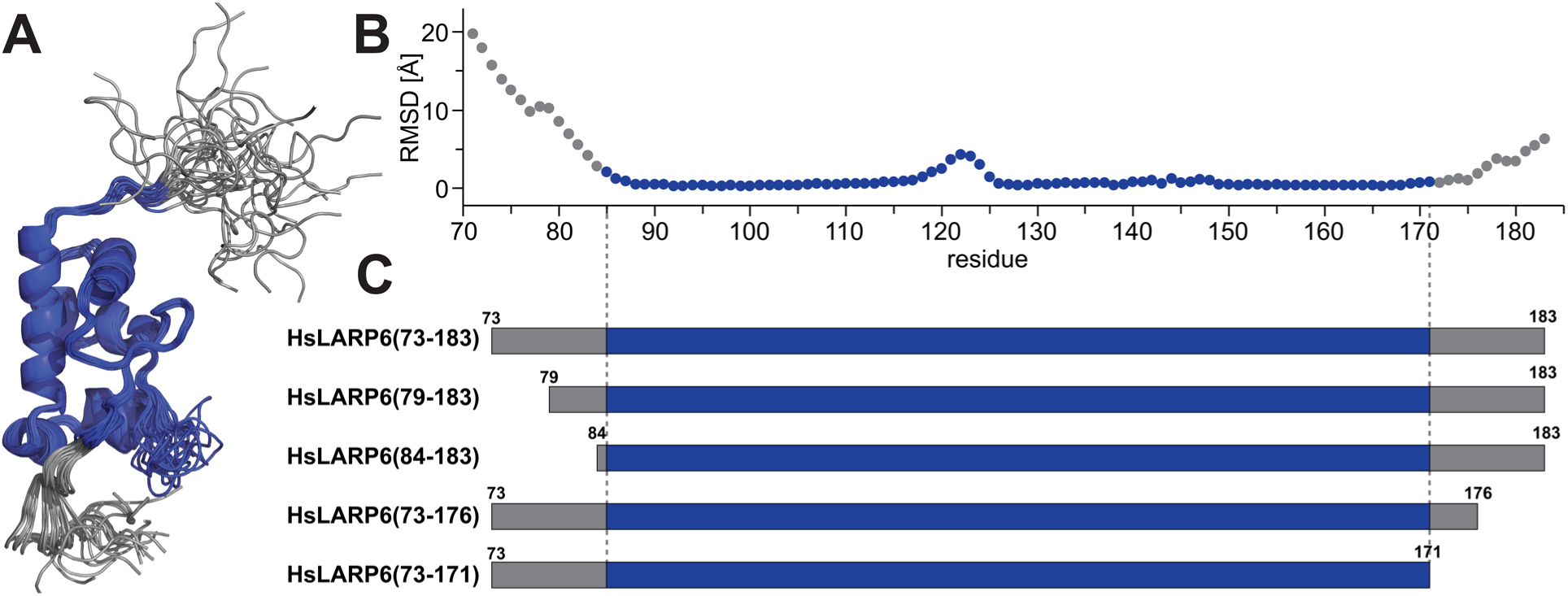
Domain boundaries of the La domain of HsLARP6. (A) The 20 lowest energy conformers of the solution NMR structure of the La domain of HsLARP6 (PDB ID: 2MTF)^27^ reveal the presence of a structured core domain (blue) with flexible regions (gray) located N- and C-terminally to the core domain. (B) Average pairwise C^α^ RMSD per residue calculated from the 20 lowest energy conformers of the NMR structure shown in (A). (C) Comparison of HsLARP6 constructs used in this study. Regions corresponding to the structured core are shown in blue and flexible N- and C-terminal extensions are shown in gray.

The most commonly studied constructs of the La domain so far span from residues T70 or G73 to S183, with only a few investigating the dependency of the length of the N- and C-terminus beyond the core domain on its ability to bind 5’SL.^16,27,31^

Hence, we decided to compare a construct spanning from residues G73 to S183, named HsLARP(73-183), to two N-terminal truncations starting at residues E79 and E84, named HsLARP(79-183) and HsLARP(84-183), respectively (Figure 4C). All three N-terminal variants vary significantly in isoelectric point (pI) with the theoretical pI of HsLARP6(73-183), HsLARP6(79-183), and HsLARP6(84-183) calculated to be 7.1, 8.2, and 9.4, respectively. The large shift in pI is explained by the presence of a large number of acidic residues located in the N-terminal tail. Indeed six of the 12 residues from 73-84 are either glutamic acid or aspartic acid. Due to the choice of a pH of 6.5 during ion exchange chromatography, the purification protocol was not adjusted for any of these variants.

All protein constructs were recombinantly expressed and purified identically to ensure side-by-side comparison (Figure 5). Similar overexpression levels were observed for all constructs (Figure S1), however, the behavior of the N-terminal and C-terminal constructs during purification was very different. However, while the purification of all N-terminal variants was successful, the C-terminal truncations HsLARP6(73-176) and HsLARP6(73-171) did not remain soluble and aggregated immediately after elution from the first IMAC column, rendering further binding studies impossible. Indeed, it was previously shown that when La domains spanning residues 81-174 and 81-166 were expressed in HEK293 cells and whole cell extracts were used in the binding assays, no binding was observed.^31^

**Figure 5:**
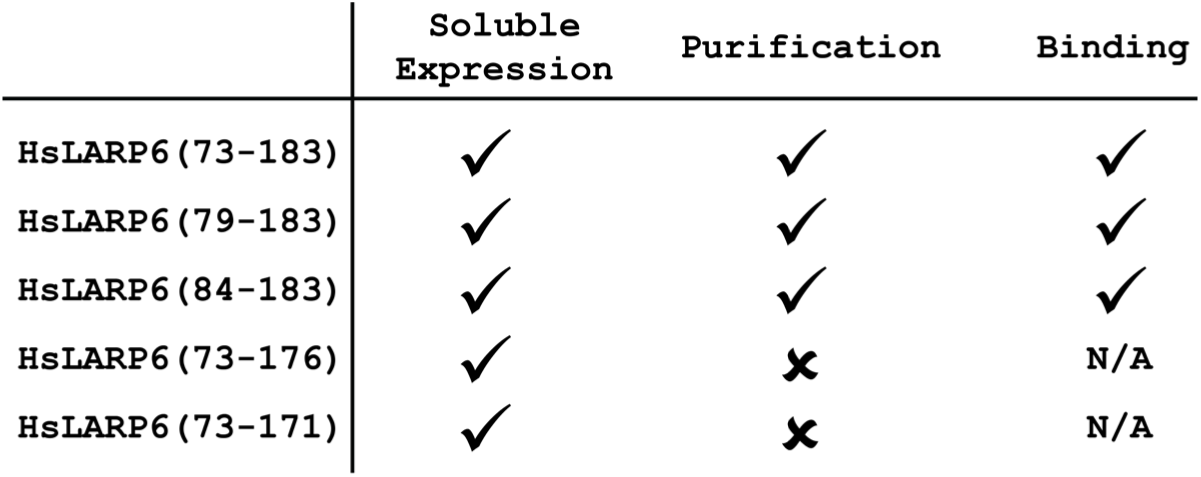
Expression and Purification of HsLARP6 La domain constructs. The HsLARP6 La domain is prone to rapid aggregation and precipitation under many common conditions. Soluble expression is possible for all constructs tested in this current study. Constructs which terminate before residue 183 did not remain in solution throughout the purification steps necessary to produce an RNAase and nucleic acid-free HsLARP6 La domain sample. Constructs which remained in solution retained binding competency.

When constructs of varying N-terminal lengths, namely HsLARP6(73-183), HsLARP6(79-183), and HsLARP6(84-183), were purified, a similar degree of stability was observed. All three constructs bind 5’SL RNA and form stable 1:1 complexes in size exclusion chromatography assays (Figure 6). All samples of the unbound state (Figure 6, black/gray traces) elute at ~ 13 ml, while chromatographs of samples of the bound state (Figure 6, red/salmon traces) show a characteristic shift to an elution volume of ~11 ml. The N-terminal variants HsLARP6(73-183), HsLARP6(79- 183), and HsLARP6(84-183) investigated here show identical binding, indicating that the N-terminal residues prior to W85 do not contribute to 5’SL binding.

**Figure 6:**
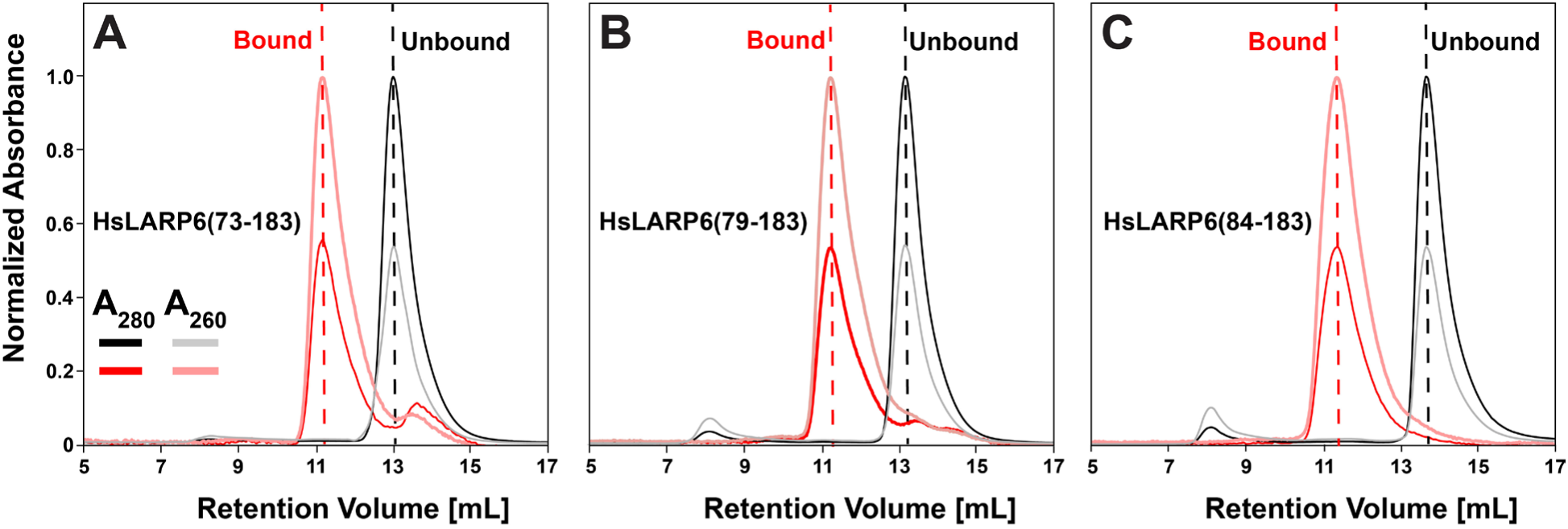
Size exclusion chromatography of 5’SL RNA-bound and unbound La domain N-terminal truncations. Overlay of the SEC profile of the normalized A_280_ (black/red) and A_260_ (gray/light red) traces of HsLARP6 residues(A) 73-183 (B) 79-183 (C) 84-183 before (black dash) and after (red dash) addition of an equimolar amount of 5’SL RNA.

### 5’SL binding stabilizes the La domain

The instability of the La domain and La module of HsLARP6 and tendency to aggregate and form precipitates is well documented.^31,33^ While observed over a large range of concentrations, the La domain of HsLARP6 is particularly aggregation-prone at higher concentrations above ~100 μM (~1.25 mg/mL), limiting further *in vitro* biochemical and biophysical studies.^27,31,33^ Hence, previous NMR spectroscopic studies of the unbound La domain and La module were conducted in the presence of 50 mM L-glutamate and 50 mM L-arginine to enhance solubility and slow down precipitation, with limited success.^27^

We investigated the intrinsic instability of the unbound state of HsLARP6(79-183) as well as bound state employing NMR spectroscopy. We acquired [^1^H, ^15^N]-HSQC spectra of HsLARP6(79-183) (unbound state) and a 1:1 complex of HsLARP6(79-183) and 5’SL (bound state) over the course of 3 hours and 7 days, respectively (Figure 7). The use of [^1^H, ^15^N]-HSQC spectra allow for the detection of backbone and side chain amide resonances. Changes in resonance position, known as chemical shift perturbations (CSPs) indicate localized changes in chemical environment, while changes in signal intensity suggest sample loss due to precipitation, as the resonances of the precipitated species are broadened beyond the limit of detection. Resonance positions and intensities of [^1^H, ^15^N]-HSQC spectra were compared at identical contour levels and scaling to detect any local or global structural changes in the La domain over time. Projections of a representative resonance were selected to show the change in intensity over time (Figure 7, insets).

**Figure 7:**
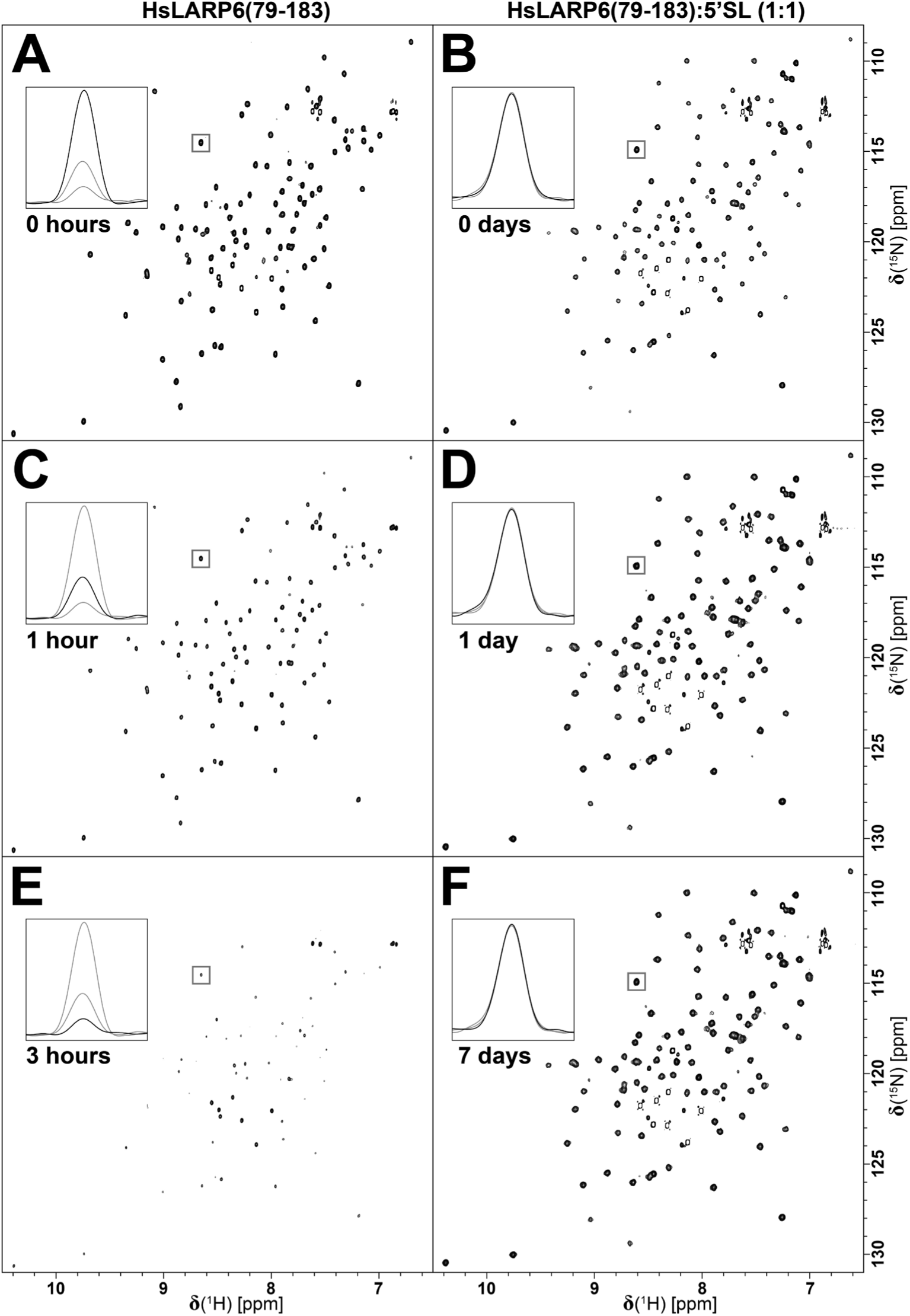
Stability of HsLAPR6(79-183) in the unbound and bound state. Signal position and intensity in 2D [^1^H, ^15^N]-HSQC, collected periodically, were employed to monitor changes in HsLARP6 La domain. The HsLARP6(79-183), in the absence of 5’SL RNA, remains soluble at 500 μM after reaching 25°C (A) but rapidly precipitates losing nearly 70% of detectable protein in 1h (B). After 3 h, the remaining signal is only ~10% of the initial intensity (C). HsLARP6(79-183) bound to 5’SL RNA is far more stable. Signal intensity at 0 h (D) remains constant after 24 h (E) and even up to 7 days (F). Insets show the 1D ^1^H proton projection of the same peak (gray box). Spectra were collected at 298 K and 16.4 T and processed identically and scaled to the same contour level and contour increment.

The unbound La domain was poorly stable as evidenced by the dramatic loss in signal intensity within the first three hours of measurement at 25°C (Figure 7, left column), as expected. Compared to a [^1^H, ^15^N]-HSQC spectrum of a freshly prepared sample of 500 μM HsLARP6(79-183) (Figure 7A), peak intensities decreased by 70% within 1 hour (Figure 7C). Peak intensities decreased by 90% within 3 hours (Figure 7E) compared to the intensity of the original spectrum. The resonances observed over the course of these measurements do not show chemical shift perturbations and no other resonances are observed, indicating that the loss in signal can be attributed to the loss of the monomeric protein species without forming any appreciable amounts of soluble aggregates. Signal loss can therefore be attributed to the formation of large (solid) protein aggregates that do not present with resonances in solution NMR spectroscopic experiments. This is further evidenced by the emergence of visible precipitation after only ~4 hours of data acquisition (Figure S2). While the process of precipitation is naturally accelerated by temperature, sample concentration, and buffer conditions, these results clearly showcased the intrinsic instability of the La domain.

Surprisingly, the binding of 5’SL RNA to the La domain of HsLARP6 counteracts this intrinsic instability and suspends precipitation entirely under otherwise identical conditions. Following the same experimental setup, the 1:1 complex of HsLARP6(79-183) and 5’SL (bound state) was stable at 25°C over the course of 7 days (Figure 7, right column). Compared to a [^1^H, ^15^N]-HSQC spectrum of a freshly prepared sample of 500 μM HsLARP6(79-183):5’SL (Figure 7B), peak intensities in [^1^H, ^15^N]-HSQC spectra recorded after 1 day and 7 days remained stable (Figure 7D & F, respectively). The [^1^H, ^15^N]-HSQC spectra recorded over the course of 7 days show no change in peak intensity or position and no visible precipitation was observed. The sample was stored at 25°C between experiments. It is worth mentioning that in our experience, various NMR samples of the HsLARP6(79-183):5’SL complex ranging from concentrations of 250 μM to 1.7 mM were measured extensively under identical conditions at 25°C, but stored at 4°C between measurements, remained stable with unchanged NMR spectra over at least 3 months. This demonstrates the tremendous shift in stability from aggregation-prone to aggregation-resistant of the bound La domain compared to the unbound La domain.

The exact source of this significant shift in stability remains elusive, however, the fact that almost all residues experience chemical shift perturbations upon RNA binding (Figure 3C) offers some insights. While it is known that the buried surface area of protein-RNA complexes is larger, on average, compared to protein-protein interactions,^34^ it is still most typical to only see CSPs at and around the binding interface.^35,36^ The presence of CSPs for nearly all residues of the bound-state protein is indicative of a global shift in conformational dynamics upon RNA binding. While the magnitude of CSPs does not indicate a completely different fold of the protein, the binding of the RNA imposes a global effect on residues far beyond the binding interface, likely stabilizing the protein fold and thus suppressing the formation of aggregation-prone species.

## Conclusions

This study details an improved purification scheme for producing milligram quantities of RNase-free, binding competent HsLARP6. The use of La domain constructs prepared freshly as proposed here showed great reproducibility. Furthermore, we showed that while variation in the length of the N-terminus between residues 73 to 84 heavily modulates the pI of the protein significantly, the expression, purification, and binding was not affected, indicating the flexible N-terminal region of the La domain is not involved in 5’SL binding. Removal of residues from the C-terminus before residue 183 greatly destabilizes the HsLARP6 La domain. While these constructs showed soluble recombinant expression, they immediately aggregated after IMAC purification. Finally, the stability of the La domain is significantly increased when bound to 5’SL, even at very high concentrations. The ribonucleoprotein complex formed is identical in NMR spectra for 7 days at room temperature while the unbound La domain in comparison shows ~90% loss in signal intensity within 3 hours under otherwise identical conditions, showcasing an incredible shift from aggregation-prone to aggregation-resistant behavior upon 5’SL binding.

## Materials and Methods

### Constructs

Sequences for various HsLARP6 La domain constructs (UniProtKB accession number Q9BRS8, Table S1) named His_6_-TEV-HsLARP6(73-183), His_6_-TEV-HsLARP6(79-183), His_6_-TEV-HsLARP6(84-183), His_6_-TEV-HsLARP6(73-176), and His_6_-TEV-HsLARP6(73-171) as well as a cysteine point mutation at position 74 named His_6_-HsLARP(73-183)G74C were cloned into a pET28a vectors via *Nco*I and *Bam*HI restriction sites for expression in *E. coli* (Genscript USA Inc.). A tobacco etch virus (TEV) protease cleavage site (ENLYFQ^▾^G) was inserted between a six histidine fusion tag (His_6_ tag) and the first residue of the native sequence.

### Chemicals

All reagents and chemicals were purchased from commercial suppliers and used as obtained. For a complete list of chemicals and their abbreviations, see Table S2.

### Expression of HsLARP6 La Domain

The plasmids coding for various His_6_-TEV-HsLARP6 La domain constructs was transformed into chemically competent *E. coli* Rosetta II (DE3) (Sigma-Aldrich) for recombinant overexpression. Transformants were plated on LB-agar supplemented with 20 g/L D-glucose, 40 ug/mL kanamycin monosulfate, and 60 ug/mL chloramphenicol and grown at 37°C overnight. For precultures, 250 ml Erlenmeyer flasks were charged with 50 ml of lysogeny broth (LB) medium containing 10 g tryptone, 5 g yeast extract, and 10 g NaCl and autoclaved. Before use, precultures were supplemented with 10 g/L D-glucose, 40 ug/mL kanamycin monosulfate, and 60 ug/mL chloramphenicol. Precultures were inoculated with colonies from freshly transformed plates and grown overnight at 37°C and 220 rpm in a MaxQ™ 8000 orbital shaker (Thermo Scientific, USA). For main cultures, 2.8 L baffled Fernbach flasks were charged with 1 L of LB medium containing 10 g tryptone, 5 g yeast extract, and 10 g NaCl and autoclaved. Before use, precultures were supplemented with 10 g/L D-glucose, 40 ug/mL kanamycin monosulfate, and 60 ug/mL chloramphenicol. Main cultures were inoculated from precultures to an OD_600_ of ~0.1. Cultures were incubated at 37°C and 220 rpm until an OD_600_ of 0.6-0.8 was reached. The cultures were subsequently cold-shocked by transferring them into an ice-water bath for 30 min. Recombinant expression was induced by addition of IPTG to a final concentration of 1 mM and was allowed to continue at 18°C and 220 rpm in an MaxQ™ 8000 orbital shaker for 16 h. Cells were collected by centrifugation at 4,000 × g and 4°C for 30 min using a Beckman Coulter™ Avanti™ J-20 XPI centrifuge equipped with a JLA-8.1000 rotor before discarding the supernatant. Cell pellets were subsequently resuspended in lysis buffer containing 20 mM Tris-HCl pH 7.4, 200 mM NaCl, 50 mM monosodium L-glutamate, 50 mM of L-arginine, 0.1% w/v CHAPS, 70 mM imidazole, and 5% v/v glycerol. A protease inhibitor mix was added to the cell suspension at final concentrations of 500 μM AEBSF, 1 μM E-64, 1 μM leupeptin, and 150 nM aprotinin before they were either flash frozen in LN_2_ and stored at −20°C or immediately subjected to purification. All buffers and solutions used throughout all purifications were formulated in 0.1% v/v DEPC-treated ultrapure water. For His_6_-TEV-HsLARP6(73-183)G74C, all buffers contained 5 mM 2-mercaptoethanol.

### Purification of HsLARP6

Cells were lysed by running the cell suspension through a Microfluidizer® M-110L (Microfluidics) in 3-5 cycles at a pressure of 50 PSI while continuously cooling the lysate to 4°C using a water-ice bath. The lysate was clarified by centrifugation at 37,500 × g and 4°C using a Beckman Coulter™ Avanti™ J-20 XPI centrifuge equipped with a JA-25.50 rotor. Subsequently, the supernatant was filtered through a 0.8 μm polyethersulfone (PES) membrane (Millipore-Sigma). Immobilized metal affinity chromatography (IMAC) was performed on a ÄKTA Start™ instrument (Cytiva, USA) monitoring absorbance at 280 nm. Clarified cell lysate was subsequently purified by immobilized metal affinity chromatography (IMAC); applied at a flow rate of 5 ml/min to a 5 mL HisTrap™ HP column (Cytiva, USA) previously equilibrated with HisTrap buffer A containing 20 mM Tris-HCl pH 7.4, 200 mM NaCl, 50 mM monosodium L-glutamate, 50 mM of L-arginine, 0.1% w/v CHAPS, and 70 mM imidazole. The column was then washed at a flow rate of 5 ml/min with HisTrap buffer A until the absorbance at 280 nm reached a stable baseline. Furthermore, the column rigorously washed at a flow rate of 5 ml/min with a high-salt wash buffer containing 20 mM Tris-HCl pH 7.4, 1 M NaCl, 50 mM monosodium L-glutamate, 50 mM of L-arginine, 0.1% w/v CHAPS, and 70 mM imidazole until the absorbance at 280 nm reached a stable baseline. The column was returned to the low-salt condition by washing the column with HisTrap A buffer until conductivity reached a stable baseline. Subsequently, His_6_-TEV-HsLARP6(79-183) was eluted in a one-step gradient using a HisTrap B buffer containing 20 mM Tris-HCl pH 7.4, 200 mM NaCl, 50 mM monosodium L-glutamate, 50 mM of L-arginine, 0.1% w/v CHAPS, and 500 mM imidazole. His_6_-tag removal by TEV digestion ^37^ was performed by adding 1 mL of 1 mg/mL in-house preparations of His_6_-TEV protease ^38^ and placing it in a Spectra/Por® 3 dialysis membrane with a molecular weight cutoff (MWCO) of 3.5 kDa. The sample was dialyzed against 1 L of dialysis buffer containing 20 mM Tris-HCl pH 7.4, 200 mM NaCl, 50 mM monosodium L-glutamate, 50 mM L-arginine, 0.1% w/v CHAPS, 70 mM imidazole, and 5 mM 2-mercaptoethanol for 16 hrs. The His_6_-tag, His_6_-tagged TEV protease and other impurities were removed by reverse HisTrap purification. The digested sample was applied at a flow rate of 5 ml/min to a 5 mL HisTrap™ HP column (Cytiva, USA) previously equilibrated with HisTrap A buffer. HsLARP6(79-183) was collected in the flow-through and wash fractions. The target protein was further purified by cation exchange chromatography using a 5 mL HiTrap™ SP Fast Flow column (Cytiva, USA) previously equilibrated with SP buffer A containing 20 mM MES pH 6.5, 50 mM monosodium L-glutamate, 50 mM of L-arginine, and 0.1% w/v CHAPS. The column was then washed at a flow rate of 5 ml/min with SP buffer A until the absorbance at 280 nm reached a stable baseline. Subsequently, HsLARP6(79-183) was eluted in a one-step gradient using a SP buffer B containing 20 mM MES pH 6.5, 1 M NaCl, 50 mM monosodium L-glutamate, 50 mM of L-arginine, and 0.1% w/v CHAPS. HsLARP6 La domain was further purified and buffer exchanged by size exclusion chromatography (SEC) using an ÄKTA pure™ 25 (Cytiva, USA) monitoring absorbance at 260 nm and 280 nm on a HiLoad 16/600 75pg column (Cytiva, USA). The detailed procedure depends on the intended application (*vide infra*). Protein concentration was determined by UV-VIS spectrophotometry using the extinction coefficients listed in Table S1 as determined using the ProtParam tool (ExPasy server).^39^ Protein purity was confirmed by sodium-dodecyl-sulfate polyacrylamide electrophoresis (SDS-PAGE) on 4-15% gradient gels (Genscript, USA) stained with 0.0025% w/v each of Coomassie® Brilliant Blue G-250 and R-250 in 10% v/v ethanol, 5% v/v glacial acetic acid in water.

### RNA Synthesis and Purification

The 48nt 5’stem-loop (5’SL) RNA (Table S3) from the mRNA of the α_2_(I) polypeptide chain of human type 1 collagen was used for all binding assays. 5’SL RNA was synthesized by *in vitro* transcription (IVT) using in-house preparations of His_6_-T7 RNA polymerase P266L. ^40–42^ IVT reactions were performed as described previously. ^35^ 500 nM T7 promoter sequence strand DNA oligonucleotides and the complementary template strand for the 5’SL RNA sequence containing 2’-methoxy modified bases at the last and penultimate residues (Integrated DNA Technologies, USA).^43^ Reactions were incubated at 37°C for 16 h, then stopped by addition of EDTA to a final concentration of 50 mM. Protein components were removed from the solution by three rounds of extraction with an equal volume of 25:24:1 phenol:chloroform:2-methylbutanol. Excess phenol in solution was removed by three rounds of extraction using a 3:1 volume ratio of diethyl ether to IVT solution. Leftover diethyl ether was removed by heating to 50°C until it had boiled off. RNA solutions were diluted with an 8M urea solution to 5mL, ~4M urea to assist in the subsequent denaturing purification step. 5’SL RNA was further purified by denaturing SEC using a Hiload 16/600 75pg size exclusion column equilibrated with 20mM Tris-HCl pH 7.2, 200 mM KCl, and 7 M urea. Fractions containing purified 5’SL RNA were pooled and diluted with isopropyl alcohol to a 1:3 ratio and supplemented with ammonium acetate pH 5.2 to a final concentration of 300 mM. Precipitation reactions were incubated at 4°C for 16 h, RNA pellets were isolated by centrifugation for 15 min at 4000 × g at 4°C in a Beckman Coulter™ Allegra™ X-15R centrifuge with SX4750 swing bucket rotor. Pellets were washed three times with 80% v/v isopropanol to remove salt, then heated to 85°C for 1 h to remove excess isopropanol resulting in a glassy solid RNA pellet. Washed pellets were dissolved in 0.1% v/v DEPC-treated water to a final concentration of 20 μM RNA and 5 mM EDTA pH 7.4. 5’SL RNA was refolded at 95°C for 1 h then transferred to 4°C until the solution was completely cooled. Refolded 5’SL RNA was exchanged into NMR or MST buffers by size exclusion chromatography (*vide infra*).

### RNase Activity Assay

Nuclease activity was performed at each purification step by exposing fluorescently labeled RNA to the fraction of interest. Pooled fractions from the cell lysate supernatant, HisTrap salt wash, HisTrap eluate, SP eluate, and SEC eluate were incubated with a final concentration of 1 μM AF488-RPS6 41mer (Table S3) for 1 h at 37°C. AF488-RPS6 41mer was synthesized following the 5’ sequence of ribosomal protein s6 mRNA (AF488-RPS6 41mer) (Integrated DNA Technologies, USA). Positive and negative controls were composed of 1 μM AF488-RPS6 41mer solutions supplemented with 0.1 mg/mL RNase A or 0.1% v/v DEPC treated water, respectively. Samples were quenched by the addition of equal volume of sample loading buffer containing 90% v/v formamide in water, 10mM Tris-HCl pH 8.0, 10mM boric acid, 0.2 mM EDTA and heating to 90°C and subsequently analyzed by denaturing PAGE on 12% w/v (19:1 Acryl:Bis-acryl) 8M urea PAGE gel run at 17.5 V/cm for 1 h. Gels were imaged on a ChemiDoc™ MP Imaging System (Bio-Rad Laboratories, Inc.) using the excitation and emission filter settings for AF488.

### Microscale Thermophoresis

Microscale thermophoresis was performed using freshly prepared His6-TEV-HsLARP6(73-183) G74C, expressed and purified as described above with the addition of 2 mM TCEP in all His-tag affinity and cation exchange buffers. Protein fractions eluted from the SP column were pooled and brought to a concentration of 100 μM. 750μL of purified His_6_-TEV-HsLARP6(73-183) G74C was incubated overnight at 4°C with a 12x molar excess of Cy5-maleimide. Excess label was removed following the labeling reaction using a Superdex 75 Increase 100/300 GL column equilibrated with MST buffer containing 20 mM Tris pH 7.25, 100 mM KCl, 0.05% v/v Triton X-100, and 5 mM DTT. Fractions containing Cy5-LARP6(73-183)G74C were pooled and the protein concentration and degree of labeling were determined by UV/Vis spectrophotometry as described previously,^44^ with protein and fluorophore extinction coefficients 16,960 M^-1^ cm^-1^ and 250,000 M^-1^ cm^-1^ with correction factor of 0.03. The degree of labeling was ~ 75%. Solutions of Cy5-LARP6(73-183)G74C were diluted to 10 nM using MST buffer. Simultaneously the solution was supplemented with 20 mg/mL BSA (New England biolabs) and 10 mg/mL yeast tRNA (invitrogen) to final concentrations of 0.4 mg/mL and 0.2 mg/mL respectively. The 48nt 5’ stem-loop (5’SL) RNA sequence used was from the α2(I) strand of type 1 collagen peptide mRNA. 5’SL RNA was refolded as described above, concentrated to ~150 μM in Vivaspin® centrifugal concentrators with MWCO of 3 kDa before exchange into MST buffer using a Superdex 75 Increase 100/300 GL column. Fractions containing RNA were pooled and brought to a concentration of ~150 μM. 5’SL RNA solutions were diluted to final concentrations of 5.0 μM, 0.4 mg/mL BSA, and 0.2 mg/mL yeast tRNA. Microscale thermophoresis was performed with a constant concentration of 10 nM Cy5-LARP6(73-183)G74C and 16 1:1 serial dilutions of 5’ SL RNA ranging from 5.0 μM-152.6 pM. Thermophoresis of the 16 capillaries were recorded with a 3 s delay period prior to 15 s of IR irradiation, set to medium intensity, and a 3 s recovery period. Thermophoresis profiles were collected for 3 replicate titrations.

For each dilution, the normalized florescence intensity (F_NORM_) was calculated by dividing the average fluorescence between 0.5 s to 1.5 s after the IR laser on-time and the average fluorescence between −1.0 s to 0.0 s before the IR laser on-time.^45–47^ Each replicate data set was then individually fit using the quadratic equation for ligand binding (Eq. 1) in Prism 10.4.1 (GraphPad Software).^48^

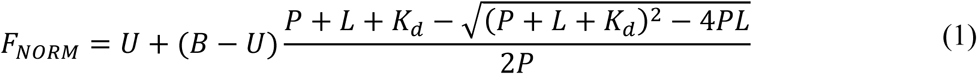

where U and B are the normalized florescence in the unbound and bound state, respectively. P is the protein concentration, L is the ligand concentration (5’SL RNA in our case), and *K_d_* is the dissociation constant.

Subsequently, U and B from individual fittings were used to first calculate the change in normalized florescence intensity (ΔF_NORM_) in by using the following relationship Δ*F_NORM_* = *F_NORM_* − *U*. The fraction bound was determined by dividing Δ*F_NORM_* by the amplitude (B-U) using the relationship *Fraction Bound* = Δ*F_NORM_*/(*B* − *U*). All three replicate datasets were then fit using the quadratic equation for ligand binding (Eq. 2) in Prism 10.4.1 (GraphPad Software).^48^

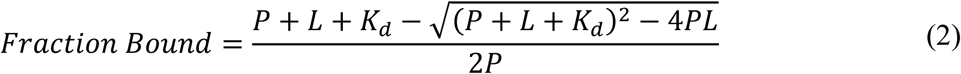

where P is the total protein concentration, L is the total ligand concentration (5’SL RNA in our case), and *K_D_* is the dissociation constant.

### Analytical size exclusion chromatography

The formation of the protein:RNA complex was explored using analytical size-exclusion chromatography (SEC). Freshly prepared protein samples of HsLARP6(73-183), HsLARP6(79-183), and HsLARP6(84-183) were purified as described above. Samples were mixed with refolded and purified 5’SL RNA in a 1:1.2 molar ratio ensuring saturation of the protein. After mixing, samples were incubated at 4°C for 10 min and subjected to analytical SEC using an AKTA pure™ 25 (Cytiva, USA) on a Superdex 75 Increase 100/300 GL column (Cytiva USA) equilibrated with 20 mM MES pH 6.5, 100 mM KCl at a flow rate of 0.75 ml/min. Absorbance at 260 nm and 280 nm was monitored.

### NMR Spectroscopy

The stability of the unbound and 5’SL RNA-bound HsLARP6 La domain was monitored using 2D [^1^H-^15^N]-HSQC experiments.^49^ Freshly prepared NMR samples of the unbound La domain [^15^N]-HsLARP6(79-183) and bound [^15^N]-HsLARP(79-183):5’SL (1:1 complex) were prepared by size exclusion chromatography (SEC) using a HiLoad 16/600 75pg column (Cytiva, USA) equilibrated with a buffer containing 11.1 mM MES pH 6.5, 55.5 mM KCl. After SEC, D_2_O and DSS were added to final concentrations of 10% v/v and 0.01% w/v, respectively, and samples were concentrated to 500 μM using Vivaspin® centrifugal concentrators with a MWCO of 3.0 kDa. All NMR data of 500 μM [^15^N]-HsLARP6(79-183) and 500 μM [^15^N]-HsLARP(79-183):5’SL (1:1 complex) in buffer containing 10 mM MES, 50 mM KCl, 10% v/v D_2_O, and 0.01% w/v DSS in 5 mm thin-walled Shigemi BMS-005B NMR tubes (Shigemi Co., Japan) were acquired at 298 K at 16.4 T on a Bruker Avance III NMR spectrometer equipped with a 3 channel ^1^H-^13^C/^15^N-^2^H, triple resonance TCI cryoprobe. 2D [^1^H, ^15^N]-HSQC experiments were recorded with 2048 by 256 complex points and 16 scans per transient. Data was directly referenced to DSS for ^1^H, while ^15^N dimensions were referenced indirectly. ^50^

Samples of both the unbound and bound state were stored at 25°C throughout the entirety of the assay. [^1^H-^15^N]-HSQC spectra of the unbound HsLARP6(79-183) at 25°C were recorded at 0 h, 1 h, and 3 h, while [^1^H-^15^N]-HSQC spectra of the 5’SL-bound HsLARP6(79-183) were recorded at 0 h, 1 day, and 7 days.

## Acknowledgements

Research reported in this publication was supported by NIGMS of the National Institutes of Health under award number R35GM142912. The content is solely the responsibility of the authors and does not necessarily represent the official views of the National Institutes of Health. The authors would like to thank Branko Stefanovic for fruitful discussion.

## Accession Codes

Sequence of Human LARP6: Uniprot Q9BRS8-1

## References

(1) Bousquet-Antonelli, C.; Deragon, J. M. A comprehensive analysis of the La-motif protein superfamily. RNA 2009, 15 (5), 750–764. DOI: 10.1261/rna.1478709 From NLM Medline.

(2) Deragon, J. M. Distribution, organization an evolutionary history of La and LARPs in eukaryotes. RNA Biol 2021, 18 (2), 159–167. DOI: 10.1080/15476286.2020.1739930 From NLM Medline.

(3) Sommer, G.; Heise, T. Role of the RNA-binding protein La in cancer pathobiology. RNA Biology 2021, 18 (2), 218–236. DOI: 10.1080/15476286.2020.1792677.

(4) Stavraka, C.; Blagden, S. The La-Related Proteins, a Family with Connections to Cancer. Biomolecules 2015, 5 (4), 2701–2722.

(5) Xie, C.; Huang, L.; Xie, S.; Xie, D.; Zhang, G.; Wang, P.; Peng, L.; Gao, Z. LARP1 predict the prognosis for early-stage and AFP-normal hepatocellular carcinoma. J Transl Med 2013, 11, 272. DOI: 10.1186/1479-5876-11-272.

(6) Zou, T.; Wang, P. L.; Gao, Y.; Liang, W. T. Circular RNA_LARP4 is lower expressed and serves as a potential biomarker of ovarian cancer prognosis. Eur Rev Med Pharmacol Sci 2018, 22 (21), 7178–7182. DOI: 10.26355/eurrev_201811_16250.

(7) Parsons, C. J.; Stefanovic, B.; Seki, E.; Aoyama, T.; Latour, A. M.; Marzluff, W. F.; Rippe, R. A.; Brenner, D. A. Mutation of the 5’ untranslated region stem-loop structure inhibits {alpha}1(i) collagen expression in vivo. J Biol Chem 2011, 286 (10), 8609–8619. DOI: 10.1074/jbc.M110.189118.

(8) Stefanovic, B.; Stefanovic, L.; Manojlovic, Z. Imaging of type I procollagen biosynthesis in cells reveals biogenesis in highly organized bodies; Collagenosomes. Matrix Biol Plus 2021, 12, 100076. DOI: 10.1016/j.mbplus.2021.100076.

(9) Makarev, E.; Izumchenko, E.; Aihara, F.; Wysocki, P. T.; Zhu, Q.; Buzdin, A.; Sidransky, D.; Zhavoronkov, A.; Atala, A. Common pathway signature in lung and liver fibrosis. Cell Cycle 2016, 15 (13), 1667–1673. DOI: 10.1080/15384101.2016.1152435.

(10) Karsdal, M. A.; Manon-Jensen, T.; Genovese, F.; Kristensen, J. H.; Nielsen, M. J.; Sand, J. M.; Hansen, N. U.; Bay-Jensen, A. C.; Bager, C. L.; Krag, A.;, et al. Novel insights into the function and dynamics of extracellular matrix in liver fibrosis. Am J Physiol Gastrointest Liver Physiol 2015, 308 (10), G807–830. DOI: 10.1152/ajpgi.00447.2014.

(11) Zhang, Y.; Stefanovic, B. Akt mediated phosphorylation of LARP6; critical step in biosynthesis of type I collagen. Scientific reports 2016, 6, 22597. DOI: 10.1038/srep22597.

(12) Wang, H.; Stefanovic, B. Role of LARP6 and Nonmuscle Myosin in Partitioning of Collagen mRNAs to the ER Membrane. PLoS One 2014, 9 (10), e108870.

(13) Blackstock, C. D.; Higashi, Y.; Sukhanov, S.; Shai, S. Y.; Stefanovic, B.; Tabony, A. M.; Yoshida, T.; Delafontaine, P. Insulin-like growth factor-1 increases synthesis of collagen type I via induction of the mRNA-binding protein LARP6 expression and binding to the 5’ stem-loop of COL1a1 and COL1a2 mRNA. J Biol Chem 2014, 289 (11), 7264–7274. DOI: 10.1074/jbc.M113.518951.

(14) Vukmirovic, M.; Manojlovic, Z.; Stefanovic, B. Serine-threonine kinase receptor-associated protein (STRAP) regulates translation of type I collagen mRNAs. Mol Cell Biol 2013, 33 (19), 3893–3906. DOI: 10.1128/MCB.00195-13.

(15) Manojlovic, Z.; Stefanovic, B. A novel role of RNA helicase A in regulation of translation of type I collagen mRNAs. RNA 2012, 18 (2), 321–334. DOI: 10.1261/rna.030288.111.

(16) Cai, L.; Fritz, D.; Stefanovic, L.; Stefanovic, B. Binding of LARP6 to the conserved 5’ stem-loop regulates translation of mRNAs encoding type I collagen. J Mol Biol 2010, 395 (2), 309–326. DOI: 10.1016/j.jmb.2009.11.020.

(17) Zhang, Y.; Stefanovic, B. LARP6 Meets Collagen mRNA: Specific Regulation of Type I Collagen Expression. Int J Mol Sci 2016, 17 (3), 419. DOI: 10.3390/ijms17030419 From NLM Medline.

(18) Stefanovic, B.; Manojlovic, Z.; Vied, C.; Badger, C. D.; Stefanovic, L. Discovery and evaluation of inhibitor of LARP6 as specific antifibrotic compound. Scientific reports 2019, 9 (1), 326. DOI: 10.1038/s41598-018-36841-y.

(19) Stefanovic, L.; Longo, L.; Zhang, Y.; Stefanovic, B. Characterization of binding of LARP6 to the 5’ stem-loop of collagen mRNAs: implications for synthesis of type I collagen. RNA Biol 2014, 11 (11), 1386–1401. DOI: 10.1080/15476286.2014.996467 From NLM Medline.

(20) Al-Ashtal, H. A.; Rubottom, C. M.; Leeper, T. C.; Berman, A. J. The LARP1 La-Module recognizes both ends of TOP mRNAs. RNA Biology 2021, 18 (2), 248–258. DOI: 10.1080/15476286.2019.1669404.

(21) Aoki, K.; Adachi, S.; Homoto, M.; Kusano, H.; Koike, K.; Natsume, T. LARP1 specifically recognizes the 3′ terminus of poly(A) mRNA. FEBS Letters 2013, 587 (14), 2173–2178. DOI: 10.1016/j.febslet.2013.05.035.

(22) Kozlov, G.; Mattijssen, S.; Jiang, J.; Nyandwi, S.; Sprules, T.; Iben, James R.; Coon, Steven L.; Gaidamakov, S.; Noronha, A. M.; Wilds, Christopher J.; et al. Structural basis of 3′-end poly(A) RNA recognition by LARP1. Nucleic Acids Research 2022, 50 (16), 9534–9547. DOI: 10.1093/nar/gkac696 (acccessed 4/27/2024).

(23) Teplova, M.; Yuan, Y.-R.; Phan, A. T.; Malinina, L.; Ilin, S.; Teplov, A.; Patel, D. J. Structural Basis for Recognition and Sequestration of UUUOH 3′ Temini of Nascent RNA Polymerase III Transcripts by La, a Rheumatic Disease Autoantigen. Molecular Cell 2006, 21 (1), 75–85. DOI: 10.1016/j.molcel.2005.10.027.

(24) Uchikawa, E.; Natchiar, K. S.; Han, X.; Proux, F.; Roblin, P.; Zhang, E.; Durand, A.; Klaholz, B. P.; Dock-Bregeon, A.-C. Structural insight into the mechanism of stabilization of the 7SK small nuclear RNA by LARP7. Nucleic Acids Research 2015, 43 (6), 3373–3388. DOI: 10.1093/nar/gkv173 (acccessed 4/27/2024).

(25) Wang, Y.; He, Y.; Wang, Y.; Yang, Y.; Singh, M.; Eichhorn, C. D.; Cheng, X.; Jiang, Y. X.; Zhou, Z. H.; Feigon, J. Structure of LARP7 Protein p65–telomerase RNA Complex in Telomerase Revealed by Cryo-EM and NMR. Journal of Molecular Biology 2023, 435 (11), 168044. DOI: 10.1016/j.jmb.2023.168044.

(26) Kuspert, M.; Murakawa, Y.; Schaffler, K.; Vanselow, J. T.; Wolf, E.; Juranek, S.; Schlosser, A.; Landthaler, M.; Fischer, U. LARP4B is an AU-rich sequence associated factor that promotes mRNA accumulation and translation. Rna 2015, 21 (7), 1294–1305, Article. DOI: 10.1261/rna.051441.115.

(27) Martino, L.; Pennell, S.; Kelly, G.; Busi, B.; Brown, P.; Atkinson, R. A.; Salisbury, N. J.; Ooi, Z. H.; See, K. W.; Smerdon, S. J.;, et al. Synergic interplay of the La motif, RRM1 and the interdomain linker of LARP6 in the recognition of collagen mRNA expands the RNA binding repertoire of the La module. Nucleic Acids Res 2015, 43 (1), 645–660. DOI: 10.1093/nar/gku1287 From NLM Medline.

(28) Yamada, Y.; Mudryj, M.; de Crombrugghe, B. A uniquely conserved regulatory signal is found around the translation initiation site in three different collagen genes. J Biol Chem 1983, 258 (24), 14914–14919.

(29) Stefanovic, B.; Hellerbrand, C.; Brenner, D. A. Regulatory role of the conserved stem-loop structure at the 5’ end of collagen alpha1(I) mRNA. Mol Cell Biol 1999, 19 (6), 4334–4342. DOI: 10.1128/MCB.19.6.4334 From NLM Medline.

(30) Castro, J. M.; Horn, D. A.; Pu, X.; Lewis, K. A. Recombinant expression and purification of the RNA-binding LARP6 proteins from fish genetic model organisms. Protein Expr Purif 2017, 134, 147–153. DOI: 10.1016/j.pep.2017.04.004 From NLM Medline.

(31) Stefanovic, L.; Gordon, B. H.; Silvers, R.; Stefanovic, B. Characterization of Sequence-Specific Binding of LARP6 to the 5’ Stem-Loop of Type I Collagen mRNAs and Implications for Rational Design of Antifibrotic Drugs. J Mol Biol 2022, 434 (2), 167394. DOI: 10.1016/j.jmb.2021.167394 From NLM Medline.

(32) Manchester, K. L. Use of UV methods for measurement of protein and nucleic acid concentrations. Biotechniques 1996, 20 (6), 968–970. DOI: 10.2144/96206bm05 From NLM Medline.

(33) Lizarrondo, J.; Dock-Bregeon, A. C.; Martino, L.; Conte, M. R. Structural dynamics in the La-module of La-related proteins. RNA Biol 2021, 18 (2), 194–206. DOI: 10.1080/15476286.2020.1733799 From NLM Medline.

(34) Kruger, D. M.; Neubacher, S.; Grossmann, T. N. Protein-RNA interactions: structural characteristics and hotspot amino acids. RNA 2018, 24 (11), 1457–1465. DOI: 10.1261/rna.066464.118 From NLM Medline.

(35) Schnieders, R.; Knezic, B.; Zetzsche, H.; Sudakov, A.; Matzel, T.; Richter, C.; Hengesbach, M.; Schwalbe, H.; Furtig, B. NMR Spectroscopy of Large Functional RNAs: From Sample Preparation to Low-Gamma Detection. Curr Protoc Nucleic Acid Chem 2020, 82 (1), e116. DOI: 10.1002/cpnc.116 From NLM Medline.

(36) Furtig, B.; Richter, C.; Wohnert, J.; Schwalbe, H. NMR spectroscopy of RNA. Chembiochem 2003, 4 (10), 936–962. DOI: 10.1002/cbic.200300700 From NLM Medline.

(37) Raran-Kurussi, S.; Cherry, S.; Zhang, D.; Waugh, D. S. Removal of Affinity Tags with TEV Protease. Methods Mol Biol 2017, 1586, 221–230. DOI: 10.1007/978-1-4939-6887-9_14 From NLM Medline.

(38) Tropea, J. E.; Cherry, S.; Waugh, D. S. Expression and purification of soluble His(6)-tagged TEV protease. Methods Mol Biol 2009, 498, 297–307. DOI: 10.1007/978-1-59745-196-3_19 From NLM Medline.

(39) Gasteiger, E.; Hoogland, C.; Gattiker, A.; Duvaud, S. E.; Wilkins, M. R.; Appel, R. D.; Bairoch, A. Protein Identification and Analysis Tools on the ExPASy Server. In The Proteomics Protocols Handbook, Walker, J. M. Ed.; Humana Press, 2005; pp 571–607.

(40) Guillerez, J.; Lopez, P. J.; Proux, F.; Launay, H.; Dreyfus, M. A mutation in T7 RNA polymerase that facilitates promoter clearance. Proc Natl Acad Sci U S A 2005, 102 (17), 5958–5963. DOI: 10.1073/pnas.0407141102 From NLM Medline.

(41) Ramirez-Tapia, L. E.; Martin, C. T. New insights into the mechanism of initial transcription: the T7 RNA polymerase mutant P266L transitions to elongation at longer RNA lengths than wild type. J Biol Chem 2012, 287 (44), 37352–37361. DOI: 10.1074/jbc.M112.370643 From NLM Medline.

(42) Ellinger, T.; Ehricht, R. Single-step purification of T7 RNA polymerase with a 6-histidine tag. Biotechniques 1998, 24 (5), 718–720. DOI: 10.2144/98245bm03 From NLM Medline.

(43) Kao, C.; Zheng, M.; Rudisser, S. A simple and efficient method to reduce nontemplated nucleotide addition at the 3 terminus of RNAs transcribed by T7 RNA polymerase. RNA 1999, 5 (9), 1268–1272. DOI: 10.1017/s1355838299991033 From NLM Medline.

(44) Wessendorf, M. W.; Tallaksen-Greene, S. J.; Wohlhueter, R. M. A spectrophotometric method for determination of fluorophore-to-protein ratios in conjugates of the blue fluorophore 7-amino-4-methylcoumarin-3-acetic acid (AMCA). J Histochem Cytochem 1990, 38 (1), 87–94. DOI: 10.1177/38.1.1688452 From NLM Medline.

(45) Wienken, C. J.; Baaske, P.; Rothbauer, U.; Braun, D.; Duhr, S. Protein-binding assays in biological liquids using microscale thermophoresis. Nat Commun 2010, 1, 100. DOI: 10.1038/ncomms1093.

(46) Jerabek-Willemsen, M.; Wienken, C. J.; Braun, D.; Baaske, P.; Duhr, S. Molecular interaction studies using microscale thermophoresis. Assay Drug Dev Technol 2011, 9 (4), 342–353. DOI: 10.1089/adt.2011.0380.

(47) Silvers, R.; Saxena, K.; Kudlinzki, D.; Schwalbe, H. Recombinant expression and purification of human TATA binding protein using a chimeric fusion. Protein Expression and Purification 2012, 85 (1), 142–147. DOI: 10.1016/j.pep.2012.07.006.

(48) Hulme, E. C.; Trevethick, M. A. Ligand binding assays at equilibrium: validation and interpretation. Br J Pharmacol 2010, 161 (6), 1219–1237. DOI: 10.1111/j.1476-5381.2009.00604.x From NLM Medline.

(49) Lehner, F.; Kudlinzki, D.; Richter, C.; Müller-Werkmeister, H. M.; Eberl, K. B.; Bredenbeck, J.; Schwalbe, H.; Silvers, R. Impact of Azidohomoalanine Incorporation on Protein Structure and Ligand Binding. ChemBioChem 2017, 18 (23), 2340–2350. DOI: 10.1002/cbic.201700437.

(50) Harris, R. K.; Becker, E. D.; Cabral De Menezes, S. M.; Granger, P.; Hoffman, R. E.; Zilm, K. W.; International Union of, P.; Applied Chemistry, P.; Biophysical Chemistry, D. Further conventions for NMR shielding and chemical shifts IUPAC recommendations 2008. Solid State Nucl Magn Reson 2008, 33 (3), 41–56. DOI: 10.1016/j.ssnmr.2008.02.004 From NLM Medline.

