## Supporting Information for "A Robust Expression and Purification Protocol for the Production of the La Domain of Human LARP6"

### Supporting Figures

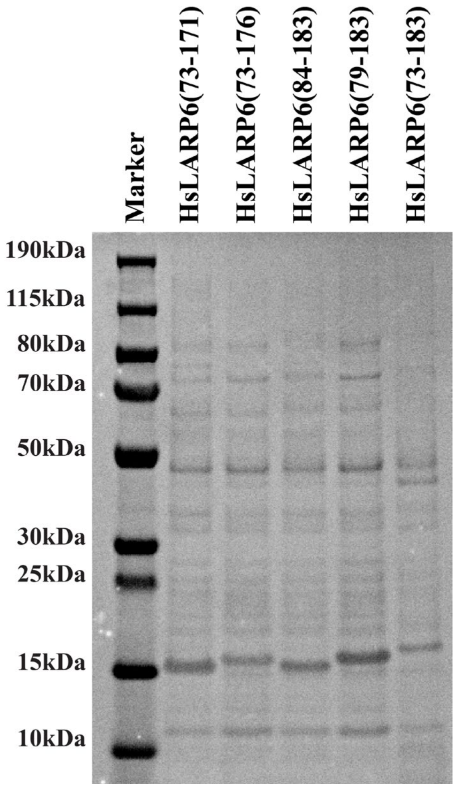


**Figure S1:** HsLARP6 La domain construct expression. SDS-PAGE gel of recombinant expressions of $His_6$-HsLARP6 residues 73-183, 79-183, 84-184, 73-176, and 73-171 (right to left) compared to PageRuler™ plus prestained protein marker. The apparent molecular weight of each band of the marker is given. 1mL of cell culture was pelleted and resuspended in 8M urea solution and heated before preparation for SDS-PAGE.

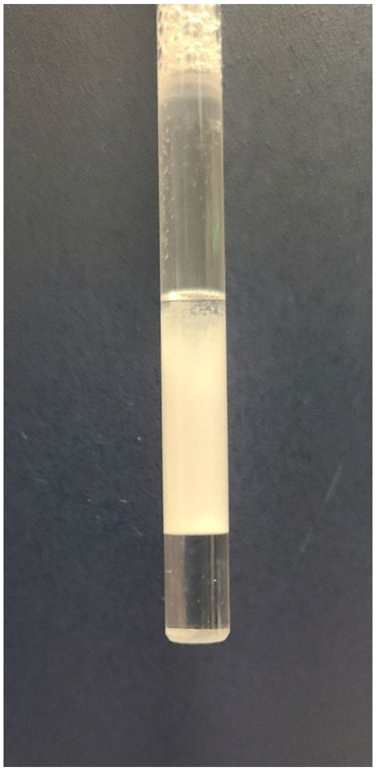

**Figure S2:** Precipitated HsLARP6(79-183). Unbound La domain photographed 4 hours after reaching 25°C for NMR measurement. The originally 500 μM La domain solution rapidly precipitated in the NMR sample tube, reaching an estimated concentration below ~ 50 μM.

```
H. sapiens               TASGGENEREDLEQEWKPPDEELIKKLVDQIEFYFSDENLEKDAFLLKHVRRNKLGYVSV
M. mulatta               TASGGENEREDLEQEWKPPDEELIKKLVDQIEFYFSDENLEKDAFLLKHVRRNKLGYVSV
B. Musculus              TTSGGENEREDLEQEWQPPDEELIKKLVDQIEFYFSDENLEKDAFLLKHVRRNKLGYVSV
F. Cats                  TTSGGENEHEDGEQEWKPPDQELIRKLVDQIEFYFSDANLEKDAFLLKHVRRNKLGYVSV
X. tropicalis            ATSGGENDGDELDQDWKPPDAELIQKLITQIEYYLSDENLEKDAFLLKHVRRNKMGFVSV
G. gallus                GRSSGGENDDDSDQNWKPPENDLIQKLVAQIEYYFSDENLEKDAFLLKHVRRNKMGYVSV
A. mississipiensis       GKSSGGENDEDFDQDWKPPENDLIQKLIEQIEYYFSDENLEKDAFLLKHVRRNKMGYVSV
                           *.* :: :: :*:*:**: :**:**: ***:*:** ****************:*:***

H. sapiens               KLLTSFKKVKHLTRDWRTTAHALKYSVVLELNEDHRKVRRTTPVPLFPNENLPSKMLLVY
M. mulatta               KLLTSFKKVKHLTRDWRTTAHALKYSVVLELNEDHRKVRRTTPVPLFPNENLPSKMLLVY
B. Musculus              KLLTSFKKVKHLTRDWRTTAHALKYSVTLELNEDHRKVRRTTPVPLFPNENLPSKMLLVY
F. Cats                  KLLTSFKKVKHLTRDWRTTAHALKYSMSLELNEDHRKVRRTTPVPLFPNENLPSKMLLVY
X. tropicalis            KLLTSFKKVKHLTRDWRTTAYALRYSNLLELNEDNRKIRRKTPVPVFPSENLPSKMLLVY
G. gallus                KLLTSFKKVKHLTRDWRTTTHALKYSDMLELNDDNRKVRRKTPVPVFPSENLPTRMLLVY
A. mississipiensis       KLLTSFKKVKHLTRDWRTTAHALKYSSILELNEDNKKVRRNTPVPVFPSENLPTRMLLVY
                         *******************::**:**  ****:*::*:**.****:**.****::*****
```

**Figure S3:** HsLARP6 protein sequence alignment performed with ClustalOmega multiple alignment tool

### Supporting Tables

**Table S1:** HsLARP6 La Domain Construct Sequences

| Construct Name | Molecular Weight [kDa] | Extinction Coefficient* [$M^{-1}cm^{-1}$] | Isoelectric point* | Sequence** |
|---|---|---|---|---|
| **His$_6$-TEV-HsLARP6(73-183)** | 15.314 (13.202) | 16960 (15470) | 6.87 (7.11) | MGHHHHHHGSGSENLYFQ\|GGENEREDLEQEWKPPDEELIKKLVDQIEFYFSDENLEKDAFLLKHVRRNKLGYVSVKLLTSFKKVKHLTRDWRTTAHALKYSVVLELNEDHRKVRRTTPVPLFPNENLPS |
| **His$_6$-TEV-HsLARP6(79-183)** | 14.728 (12.616) | 16960 (15470) | 7.34 (8.15) | MGHHHHHHGSGSENLYFQ\|GEDLEQEWKPPDEELIKKLVDQIEFYFSDENLEKDAFLLKHVRRNKLGYVSVKLLTSFKKVKHLTRDWRTTAHALKYSVVLELNEDHRKVRRTTPVPLFPNENLPS |
| **His$_6$-TEV-HsLARP6(84-183)** | 14.113 (12.002) | 16960 (15470) | 9.16 (9.40) | MGHHHHHHGSGSENLYFQ\|GEWKPPDEELIKKLVDQIEFYFSDENLEKDAFLLKHVRRNKLGYVSVKLLTSFKKVKHLTRDWRTTAHALKYSVVLELNEDHRKVRRTTPVPLFPNENLPS |
| **His$_6$-TEV-HsLARP6(73-176)** | 14.562 (12.450) | 16960 (15470) | 7.34 (8.15) | MGHHHHHHGSGSENLYFQ\|GGENEREDLEQEWKPPDEELIKKLVDQIEFYFSDENLEKDAFLLKHVRRNKLGYVSVKLLTSFKKVKHLTRDWRTTAHALKYSVVLELNEDHRKVRRTTPVPLF |
| **His$_6$-TEV-HsLARP6(73-171)** | 14.009 (11.897) | 16960 (15470) | 7.34 (8.15) | MGHHHHHHGSGSENLYFQ\|GGENEREDLEQEWKPPDEELIKKLVDQIEFYFSDENLEKDAFLLKHVRRNKLGYVSVKLLTSFKKVKHLTRDWRTTAHALKYSVVLELNEDHRKVRRTT |
| **His$_6$-TEV-HsLARP6(73-183) G74C** | 15.360 (13.248) | 16960 (15470) | 6.86 (7.11) | MGHHHHHHGSGSENLYFQ\|GCENEREDLEQEWKPPDEELIKKLVDQIEFYFSDENLEKDAFLLKHVRRNKLGYVSVKLLTSFKKVKHLTRDWRTTAHALKYSVVLELNEDHRKVRRTTPVPLFPNENLPS |

*Values in parenthesis indicate the value after TEV digestion

**Vertical line indicates the TEV cleavage site and sequence component by TEV digestion

**Table S2:** List of chemicals and their abbreviations

| Chemical name | Distributor |
|---|---|
| 2-(N-morpholino)ethanesulfonic acid (MES) | VWR |
| 2-amino-2-(hydroxymethyl)-1,3-propanediol (Tris) | VWR |
| 2-mercaptoethanol | VWR |
| 2-methylbutanol (isoamyl alcohol) | Thermo Fisher Scientific |
| 3-[(3-cholamidopropyl)dimethylammonio]-1-propanesulfonate (CHAPS) | Gold Biotechnology, Inc. |
| 40 % w/v (19:1) acrylamide:bis-acrylamide | VWR |
| 4-benzenesulfonyl fluoride hydrochloride (AEBSF) | Gold Biotechnology, Inc. |
| adenosine triphosphate | Chem-Impex International |
| ammonium carbonate | Thermo Fisher Scientific |
| ammonium chloride (15N) | Cambridge Isotope Laboratory |
| bovine pancreatic trypsin inhibitor (aprotinin) | Gold Biotechnology, Inc. |
| bovine serum albumin (BSA) | New England Biolabs |
| chloramphenicol | VWR |
| chloroform | VWR |
| Coomassie ® Brilliant Blue G-250 | VWR |
| Coomassie ® Brilliant Blue R-250 | VWR |
| cytosine triphosphate | Chem-Impex International |
| deuterium oxide (99.9%) | Cambridge Isotope Laboratory |
| D-glucose | VWR |
| diethyl dicarbonate (DEPC) | VWR |
| diethyl ether | Millipore-Sigma |
| dimethyl sulfoxide (DMSO) | Millipore-Sigma |
| dithiothreitol (DTT) | Gold Biotechnology, Inc. |
| ethanol | VWR |
| ethylenediaminetetraacetic acid (EDTA) | VWR |
| fluorescein-5-maleimide | TCI America |
| glacial acetic acid | VWR |
| glycerol | Macron Fine Chemicals |
| guanosine triphosphate | Chem-Impex International |
| imidazole | Thermo Fisher Scientific |
| isopropanol | VWR |
| isopropylthio-β-galactoside (IPTG) | Gold Biotechnology, Inc. |
| kanamycin monosulfate | Gold Biotechnology, Inc. |
| L-arginine free base | VWR |
| magnesium chloride | VWR |
| monosodium L-glutamate | TCI America |
| N-(trans-Epoxysuccinyl)-L-leucine 4-guanidinobutylamide (E-64) | Gold Biotechnology, Inc. |
| N-acetyl-L-leucyl-L-leucyl-L-arginine (leupeptin) | Gold Biotechnology, Inc. |
| phenol | Acros Organics |
| potassium chloride | Thermo Fisher Scientific |
| sodium chloride | VWR |
| sodium trimethylsilylpropanesulfonate (DSS) | Thermo Fisher Scientific |
| spermadine | TCI America |
| tris(2-carboxyethyl)phosphine hydrochloride (TCEP-HCl) | Gold Biotechnology, Inc. |
| triton x-100 | VWR |
| tryptone | Research Products International |
| uracil triphosphate | Chem-Impex International |
| urea | Acros Organics |
| yeast extract | Research Products International |
| yeast inorganic pyrophosphatase (YIPP) | New England Biolabs |

**Table S3:** RNA Sequences of the AF488-RPS6 and 5'SL RNA used in Nuclease and Stability assays

| Name | Sequence |
| --- | --- |
| **AF488-RPS6 41mer** | 5'-Alexa Fluor-488-CUC UUU UCG UGG CGC CUC GGA GGC GUU CAG CUG CUU CAA G-3' |
| **5'SL RNA (a2(I))** | 5'-GGC ACA AGG AGU CUG CAU GUC UAA GUG CUA GAC AUG CUC AGC UUU GUG-3' |
